# Does Fermentation Enhance Beverage Safety? Kombucha’s Resistance to Microbial Invasion Suggests a Protective Role

**DOI:** 10.64898/2025.12.23.696228

**Authors:** Gissel Marquez Alcaraz, Shrinath Narayanan, Joe Alcock, Jessica D. Ayers, Cristina Baciu, Hanna L. Berman, Robert R. Dunn, Angelo Fortunato, Alexander May, Carlo C. Maley, Jessica Siegel, Bryan Yavari, Athena Aktipis

**Affiliations:** Arizona Cancer Evolution Center, Biodesign Institute and School of Life Sciences, Arizona State University, Tempe, AZ, U.S.A.; Biodesign Center for Biocomputing, Security and Society, Arizona State University, Tempe, U.S.A.; Biodesign Center for Health Through Microbiomes, Arizona State University, Tempe, AZ, U.S.A.; Department of Psychological Science, Boise State University, Boise, ID, USA; Department of Psychology, Arizona State University, Tempe, AZ, U.S.A.; Department of Ecology and Evolution, University of Lausanne, Switzerland; Evolver Dynamics, Inc., Belleville, Ontario, Canada; Department of Theoretical and Applied Sciences, eCampus University, Novedrate, Italy; Department of Emergency Medicine, University of New Mexico Health Sciences Center. Albuquerque, NM, U.S.A.; Department of Applied Ecology, North Carolina State University, Raleigh, NC, USA; College of Medicine – Phoenix, University of Arizona, Phoenix, AZ, USA; Center for Evolution and Medicine, Arizona State University, Tempe, AZ, U.S.A.

**Author notes:** These authors contributed equally to this work (Gissel Marquez Alcaraz, Shrinath Narayanan). Senior author (Athena Aktipis).

## Abstract

Does fermentation enhance the safety of beverages? This study investigates kombucha’s resilience to microbial invasion from human hands. Kombucha, an ancient fermented tea, relies on a biofilm known as a symbiotic community of bacteria and yeast (SCOBY) and a starter culture to ferment sweet tea, producing various metabolites including gluconic acid, acetic acid, ethanol, and carbon dioxide. While fermented foods have been historically associated with potential health benefits, direct evidence comparing the prevalence of human pathogens in fermented versus non-fermented beverages has been lacking. To address this, we conducted a citizen science experiment at Arizona State University, introducing swabs from people’s palms into both kombucha and a sweet tea control. Over 30 days, we monitored the bacterial and fungal composition using 16S and ITS rRNA sequencing. Our findings reveal that kombucha maintained a relatively stable microbial profile and physical appearance, with a typical SCOBY biofilm developing. In stark contrast, the sweet tea experienced a dramatic change in microbial composition and was visibly compromised by rapidly growing microorganisms. Importantly, this suggests that the complex microbial ecosystem of kombucha can limit the growth of foreign microbes introduced from human hands. Given that human pathogens were absent from the kombucha while present in the tea samples, our results indicate that the antimicrobial properties of fermentation byproducts and the physical barrier of the SCOBY may contribute to this resilience. Further research is warranted to fully elucidate the mechanisms underlying kombucha’s resistance to microbial invasion.

## Introduction

Fermented foods and drinks have long provided many benefits to human societies. One of the benefits that is often mentioned is disease control. Scholars have hypothesized that beer-drinking in ancient Mesopotamia helped to reduce the exposure of people to pathogens present in contaminated drinking water ^1,2^. Similarly, it has been hypothesized that alcohol consumption among people in 19th-century Britain reduced their risk of cholera^3^. While high concentrations of alcohol can kill many pathogens, and high concentrations of alcohol are used to preserve biological specimens, it is less clear whether the biocides–be they alcohol, lactic acid, acetic acid or other compounds–have such an effect when in the concentrations at which they are consumed in fermented beverages. Indeed, some archaeologists have suggested such benefits to be unlikely in traditional beers^4^. We have been unable to find any studies that compare the prevalence or diversity of human pathogens in ferments and comparable non-fermented drinks. Here, we consider the potential of kombucha microorganisms to limit the growth of pathogens.

### Kombucha has existed for thousands of years

Kombucha is a fermented beverage believed to have originated in northeast China in 220 B.C. during the Tsin Dynasty^5^. It is one of a diversity of tea-based Chinese fermentations. One perhaps apocryphal account states that in 414 A.D., a physician named Kombu introduced the drink to Japan to treat Emperor Inkyo’s digestive issues^5,6^. Eventually, kombucha reached Russia, and from there, it continued to spread and became popular in Europe during World War II^5^. Today, kombucha is consumed globally, sold commercially, and brewed in homes. The kombucha market is worth billions of dollars and projected to continue growing^7^. The kombucha brewing methods used today have developed from an ancient and diverse tea-fermentation tradition.

### Kombucha is a multi-species ecosystem consisting of yeast and bacteria

Compared to many industrialized fermentations, which typically use single strains of single species (e.g., of *Saccharomyces cerevisiae*, known also as Brewer’s yeast), kombuchas are complex microbial ecosystems. This microbial diversity makes them similar to the kinds of natural fermentations that were brewed and consumed during human history. The fermentation process for kombucha typically takes 10–14 days. During each fermentation cycle, bacteria produce cellulose that gradually develops into a round pellicle on the surface known as the SCOBY (Symbiotic Culture of Bacteria and Yeast)^8^.

A new kombucha culture begins with “sweet tea” (tea and sugar) and a “starter”. The starter, which can be liquid and/or SCOBY from a previous batch, introduces the necessary yeast and bacteria. During fermentation, yeast cells produce invertase, which breaks down sugar (sucrose) into glucose and fructose (Figure 1). Yeasts also produce ethanol and carbon dioxide as waste products; in doing so, they contribute to the fizz and slight alcohol content. Bacteria, in turn, metabolize the glucose, fructose and ethanol to produce cellulose, forming the SCOBY, as well as acids such as acetic acid and gluconic acid, which lowers the pH^9^. In order to make an adequate kombucha, the initial starter should be 10%-20% liquid kombucha v/v (volume/volume percentage) or 2.5% SCOBY w/v (weight/volume percentage)^5^.

**Figure 1:**
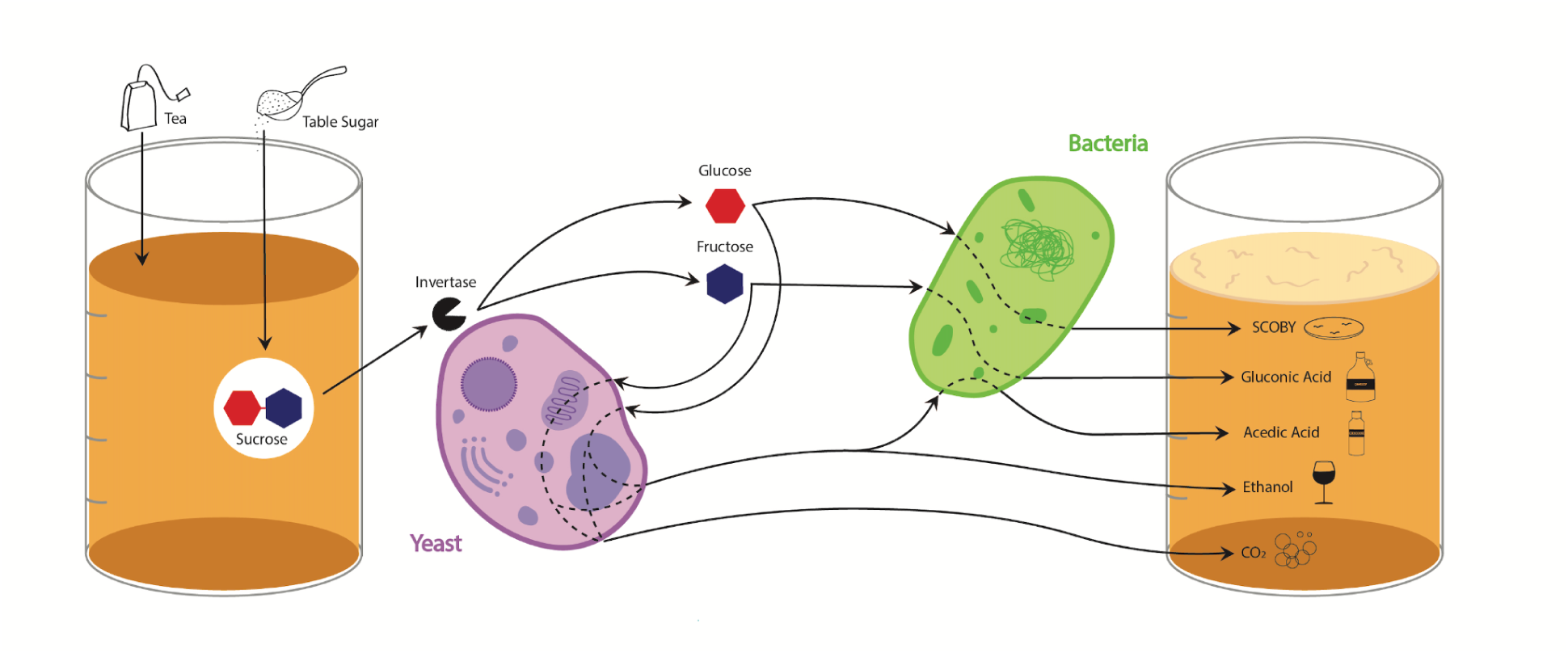
Yeasts produce an enzyme called invertase, which breaks down sucrose, allowing fermentation to begin. Yeasts also produce CO2 and Ethanol as waste. Once invertase breaks sucrose into glucose and fructose, bacteria can then metabolize the two components to create cellulose, gluconic acid, and acetic acid. Adapted from May et al. 2019^9^

Kombucha’s fermentation occurs through interactions between bacteria and yeast which produce all the potentially protective byproducts of the fermentation process, including a SCOBY, ethanol, carbon dioxide, and acids. Without yeast, sugar is not cleaved into glucose and fructose and therefore cannot be processed by bacteria to make the SCOBY and acids. Because bacteria need yeast, and each makes essential components for kombucha, their relationship is fundamentally cooperative, forming a mutualism (Table 1). The presence of the ethanol and acid by-products may be responsible for limiting the entry of pathogens from the outside environment or from the hands of a handler, a possibility that we explore in this experiment (Table 1).

**Table 1:**
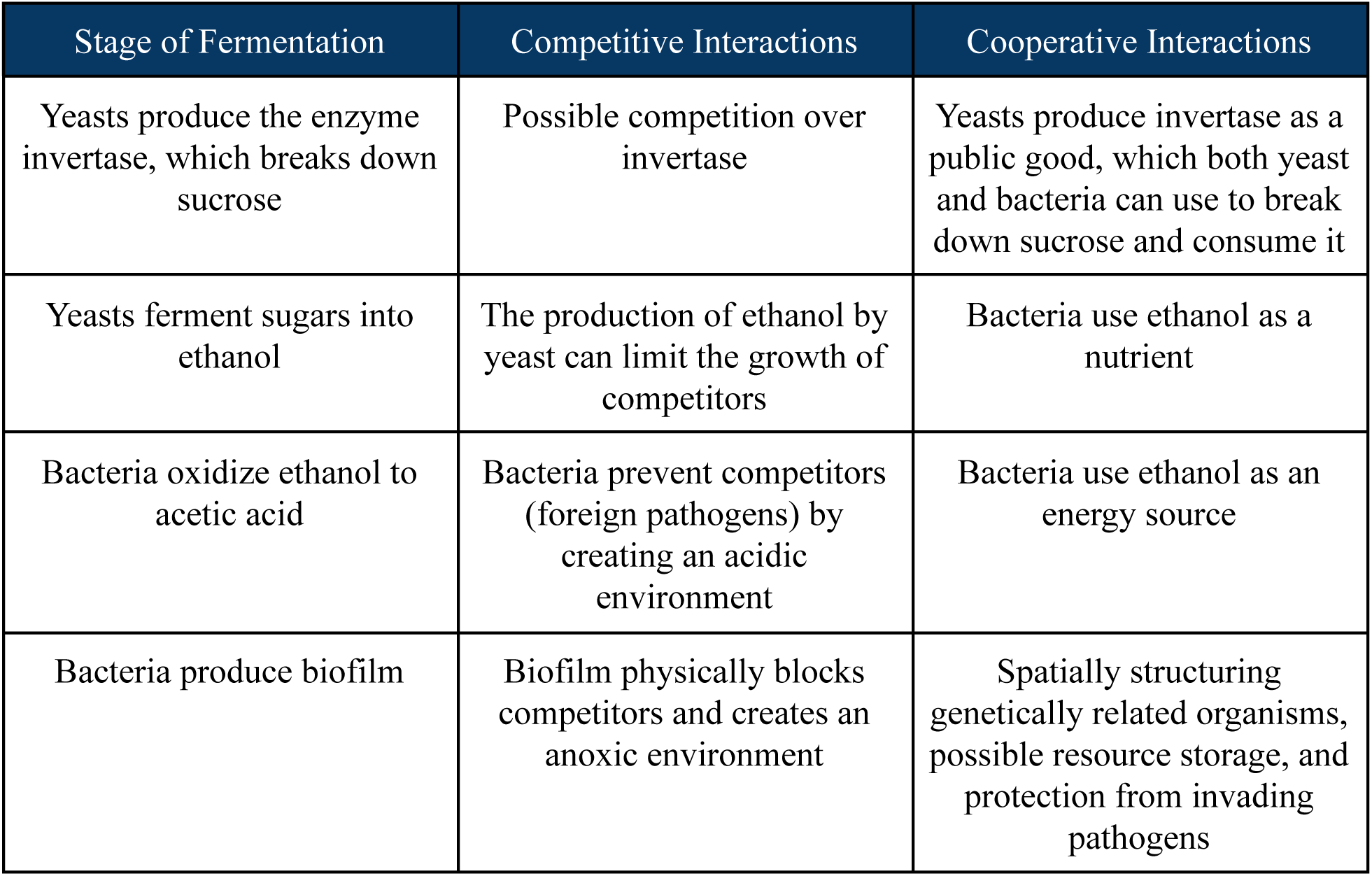
Adapted from May et al. 2019.^9^ During kombucha fermentation, microbes cooperate and compete.

### Evaluating fermentation’s potential to enhance beverage safety using kombucha

Given the historical interest in whether fermentation may contribute to the safety of beverages, and noting that kombucha possesses inherent properties such as a lower pH and a physical SCOBY barrier that might limit microbial invasion, we hypothesized that microbial fermentation would limit the growth of harmful pathogens. We designed an experiment to test whether kombucha’s established microbial ecosystem could resist the introduction of external microbes from human hands. To examine this, we introduced microbes from participants’ hands into both kombucha and sweet tea and subsequently monitored the diversity and composition of bacteria and yeast over the following 30-day period. This citizen science approach, conducted during Arizona State University’s Open Door event, involved setting up four replicates of both conditions and collecting samples on Day 1 and Day 30 for 16S rRNA and ITS sequencing and analysis. We predicted that the kombucha would have a microbial profile with fewer harmful pathogens, compared to a non-fermented sweet tea control. This would suggest that fermentation could indeed enhance beverage safety for human consumption, a question that, to our knowledge, prior research has not directly addressed.

## Results

### Kombucha SCOBYs maintained a normal appearance

The SCOBYs did not change in physical appearance between Day 1 and Day 30, whether with regard to color or other features. Between Day 1 and Day 30, the tea samples turned darker and were colonized by dark-colored microbial growth on the surface of the liquid. These changes in the tea were also observable halfway through the experiment on Day 15 (Supplementary Fig. 3).

### Bacteria increased in tea but stayed stable in kombucha; fungi declined in tea but stayed stable in kombucha

The initial mean read count of bacteria in the kombucha samples was five-fold greater (52,689) than in the tea sample (10,938) on Day 1, based on 16S data. Through time, the kombucha samples increased modestly in read count (to 61,560) (Table 2). In contrast, the increase in read count was much greater in the tea samples, with the tea samples increasing in read count to levels exceeding those of Day-1 or Day-30 SCOBYs (70,497). The use of “read counts” is, at best, an imperfect metric of absolute abundance among samples^10^. Nonetheless, the kombucha results are consistent with a model in which initial bacterial abundance and biomass were close to a stable state and where, conversely, abundance/biomass were increasing in the tea samples through time.

**Table 2:** Read counts from 16S and ITS sequencing across kombucha and tea samples at Day 1 and Day 30, showing increasing bacterial read counts over time and stable (Kombucha) or declining (Tea) fungal read counts.

|  | **16s** |  |  |  | **ITS** |  |  |  |
| --- | --- | --- | --- | --- | --- | --- | --- | --- |
|  | **Kombucha** |  | **Tea** |  | **Kombucha** |  | **Tea** |  |
| **Replicate** | **1** | **30** | **1** | **30** | 1 | 30 | 1 | 30 |
| Replicate 1 | 50839 | 73932 | 5910 | 64196 | 208532 | 194817 | 9553 | 6004 |
| Replicate 2 | 42476 | 40547 | 15204 | 63148 | 141498 | 184330 | 11695 | 1221 |
| Replicate 3 | 59634 | 58236 | 1343 | 54580 | 176038 | 218915 | 2046 | 2504 |
| Replicate 4 | 57807 | 109274 | 21294 | 64317 | 153978 | 148063 | 3917 | 2763 |

Similar to the 16S bacterial data, the number of fungal reads in kombucha was similar between Day 1 (170,011.5) and Day 30 (186,531.25) based on ITS data. In contrast to the bacterial results, tea samples showed decreases in fungal read count from Day 1 (6,802.75) to Day 30 (3,123).

### Bacterial alpha diversity declined in tea but remained stable in kombucha, while fungal alpha diversity stayed unchanged in both

#### Bacteria

At a rarefaction depth of 1343 reads, which retained all 16 samples, overall Kruskal–Wallis tests detected group-level differences in both observed Amplicon Sequence Variant (ASV) richness (H = 10.4, df = 3, p = 0.016) and Shannon index (H = 12.8, df = 3, p = 0.005) (see Supplementary Fig. 1). For ASV richness, tea showed a reduction over 30 days: Tea Day 30 had fewer ASVs than Tea Day 1 (Wilcoxon rank-sum p = 0.05), whereas kombucha ASV richness did not change. When evenness was considered with the Shannon index, Tea Day 30 showed a clear decline, and Kombucha Day 30 showed a slight increase from Kombucha Day 1 (p = 0.043 for each contrast). These findings indicate that tea lost bacterial diversity, while kombucha remained comparatively stable.

#### Fungi

Neither the ASV richness nor the Shannon index of fungi changed significantly on pairwise comparison between Day 1 and Day 30 for kombucha or tea, while there were group-level differences detected for ASV richness (H = 6.5, df =3, p = 0.089) and Shannon index (H = 11.5, df =3, p = 0.009). Fungal communities in both drinks maintained stable alpha diversity during the 30-day incubation (see Supplementary Fig. 1).

### Bacterial beta diversity changed significantly in all groups, especially in tea over time, while fungal beta diversity showed slight shifts in tea and remained stable in kombucha

#### Bacteria

For bacteria, weighted UniFrac ordination placed the four sample groups—Kombucha Day 1, Kombucha Day 30, Tea Day 1 and Tea Day 30 in distinct clusters (Figure 2). A PERMANOVA on weighted distances at a rarefaction depth of 1343 reads confirmed significant compositional differences (pseudo-F = 148.4, p = 0.001); every pairwise comparison was significant after FDR correction (q ≤ 0.035), with the greatest separation involving Tea Day 30 relative to other treatments (pseudo-F = 746–777). Repeating the analysis at 5910 reads (one sample lost) yielded the same pattern and significance. Unweighted UniFrac produced a much smaller effect (pseudo-F = 4.28, p = 0.001) and only comparisons that involved Tea Day 30 reached significance (q = 0.058). These findings indicate that changes in the relative abundance of dominant taxa drive the observed divergence among treatments. When focusing on temporal changes, tea shifted markedly over the 30 days of the experiment, whereas kombucha remained comparatively stable.

**Figure 2:**
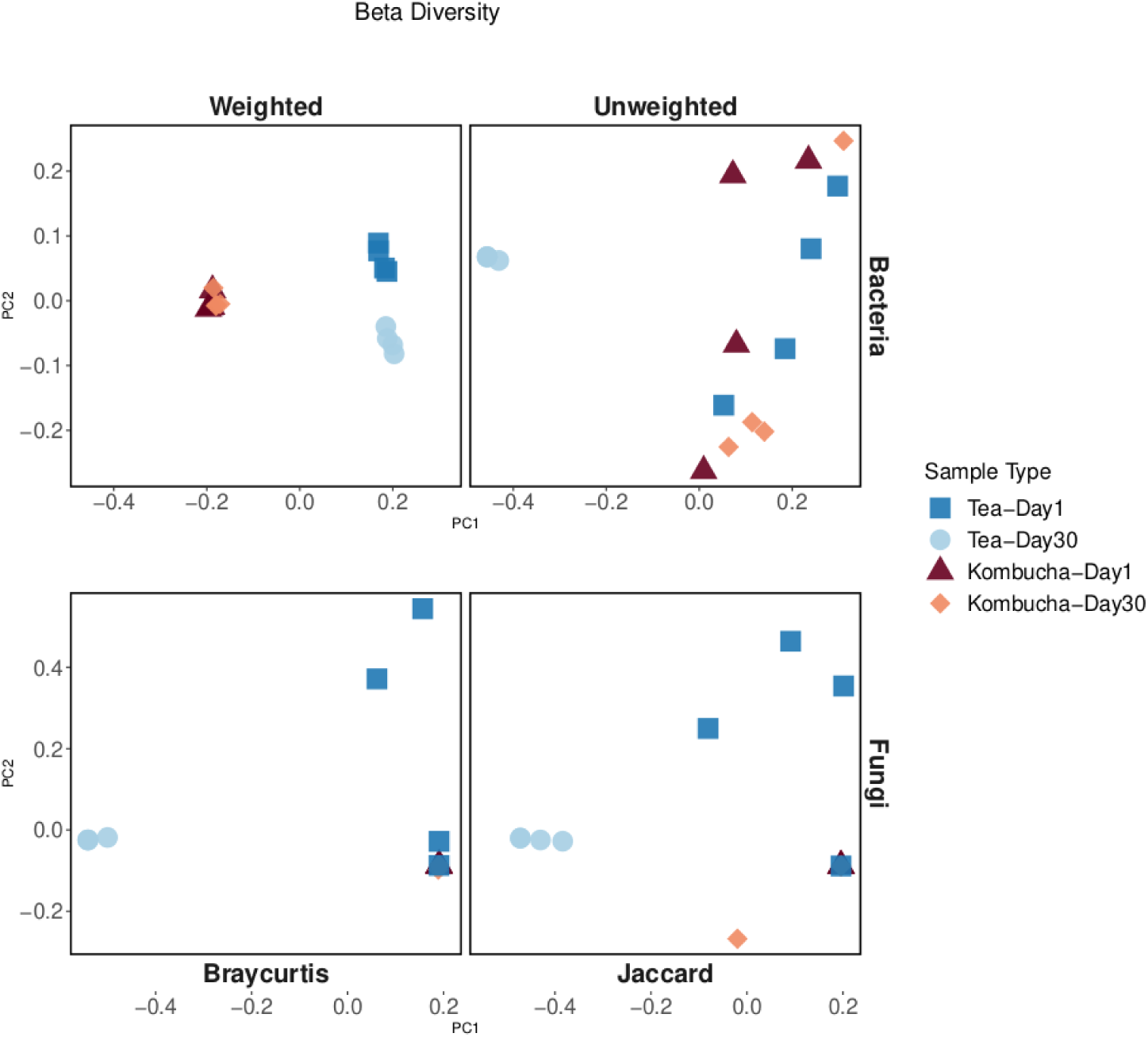
Principal coordinate analysis of bacterial (top row) and fungal (bottom row) communities in kombucha and tea samples on Day 1 and Day 30.

#### Fungi

For fungi, Bray–Curtis and Jaccard distances, rarefied to 1221 reads, also showed a significant overall group effect (pseudo-F = 2.18 and 1.67, respectively; p = 0.001 for both) (Figure 2). PCoA distinguished Tea Day 30 from the other samples, while Kombucha Day 1 and Day 30 overlapped. No pairwise comparison remained significant after FDR adjustment (minimum q = 0.062), suggesting that the fungal shift in tea is modest and that kombucha communities are temporally stable. Analyses at deeper read depths gave similar results (data not presented).

### Taxon-specific Responses: Bacterial composition changed in kombucha and tea

Between Day 1 and Day 30, bacterial diversity increased in the kombucha samples (Figure 3B), and decreased in tea samples (Figure 3A). *Komagataeibacter* remained the dominant genus of bacteria in the kombucha samples, though other genera increased in abundance, and diversity increased from Day 1 to Day 30. In the tea samples, *Burkholderia* was the most abundant on both Day 1 and Day 30. There was a drastic decrease in bacterial diversity in tea (Figure 3A).

**Figures 3A and 3B:**
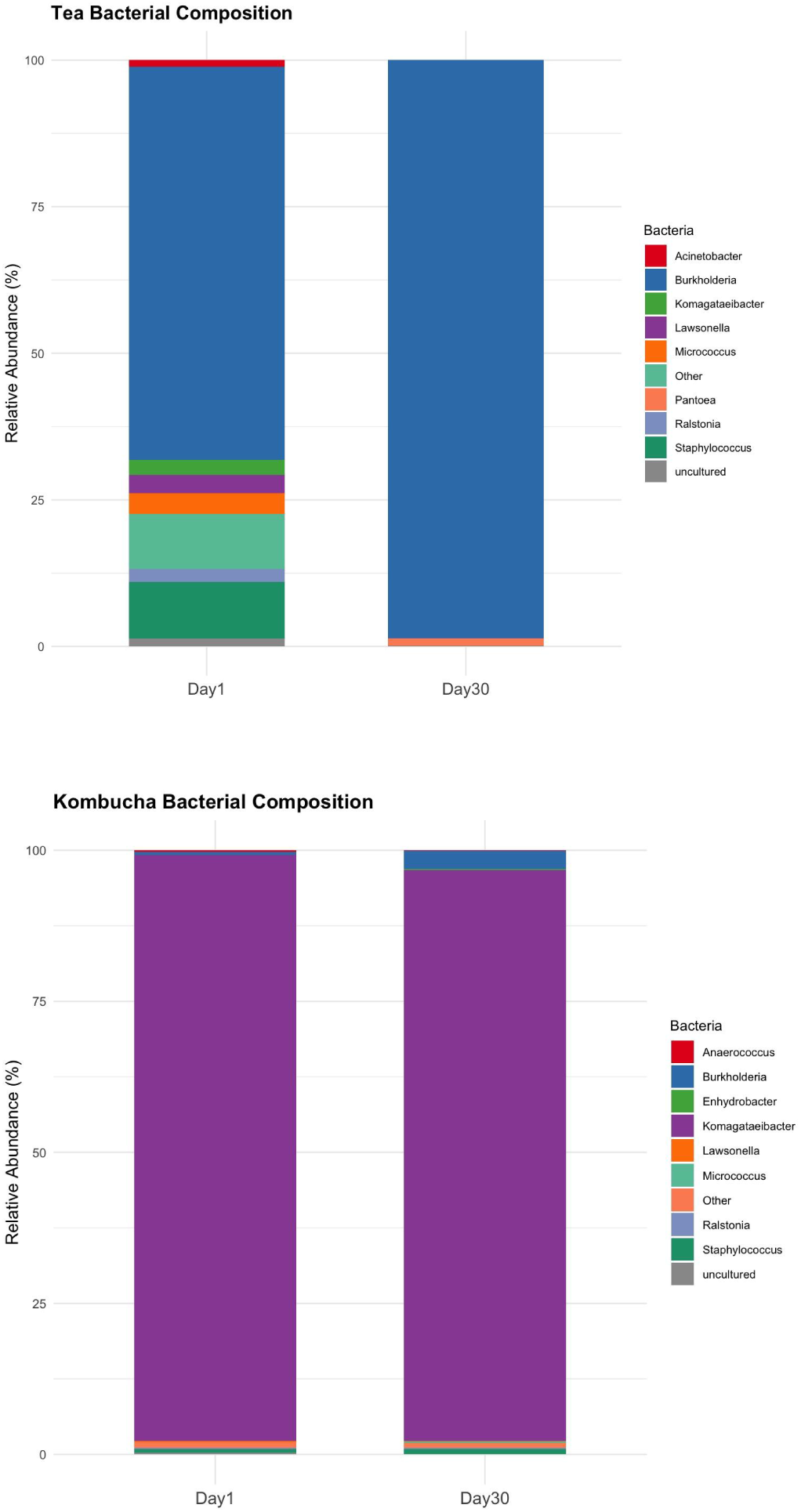
These graphs represent the mean relative abundance across all replicates. By Day 30, *Burkholderia* was the most dominant genus across tea samples. In contrast, kombucha exhibited a more stable bacterial community, with *Komagateibacter* remaining the most abundant genus throughout the study, representing 96.7% on Day 1 and 93.8% on Day 30. While there were minor changes, including an increase in *Burkholderia*, these changes were less than 5% of the overall abundance, indicating compositional stability.

### Taxon-specific Responses: Fungal composition remained stable in kombucha but changed in tea

Kombucha’s fungal profiles did not change between Day 1 and Day 30 in terms of diversity and abundance (Figure 4B). *Dekkera* persisted as the only genus identified in the kombucha replicates. These *Dekkera* species were *Dekkera bruxellensis* (most abundant) and *Dekkera anomala* (very minimal). The tea samples from Day 1 and Day 30 differed in fungal diversity and fungal abundance (Figure 4A). *Dekkera bruxellensis* was prominent in Day 1 tea, but by Day 30 *Candida hyderabadsis* was the most abundant fungi, followed by *Alternaria* and *Aureobasidium*.

**Figure 4A and 4B:**
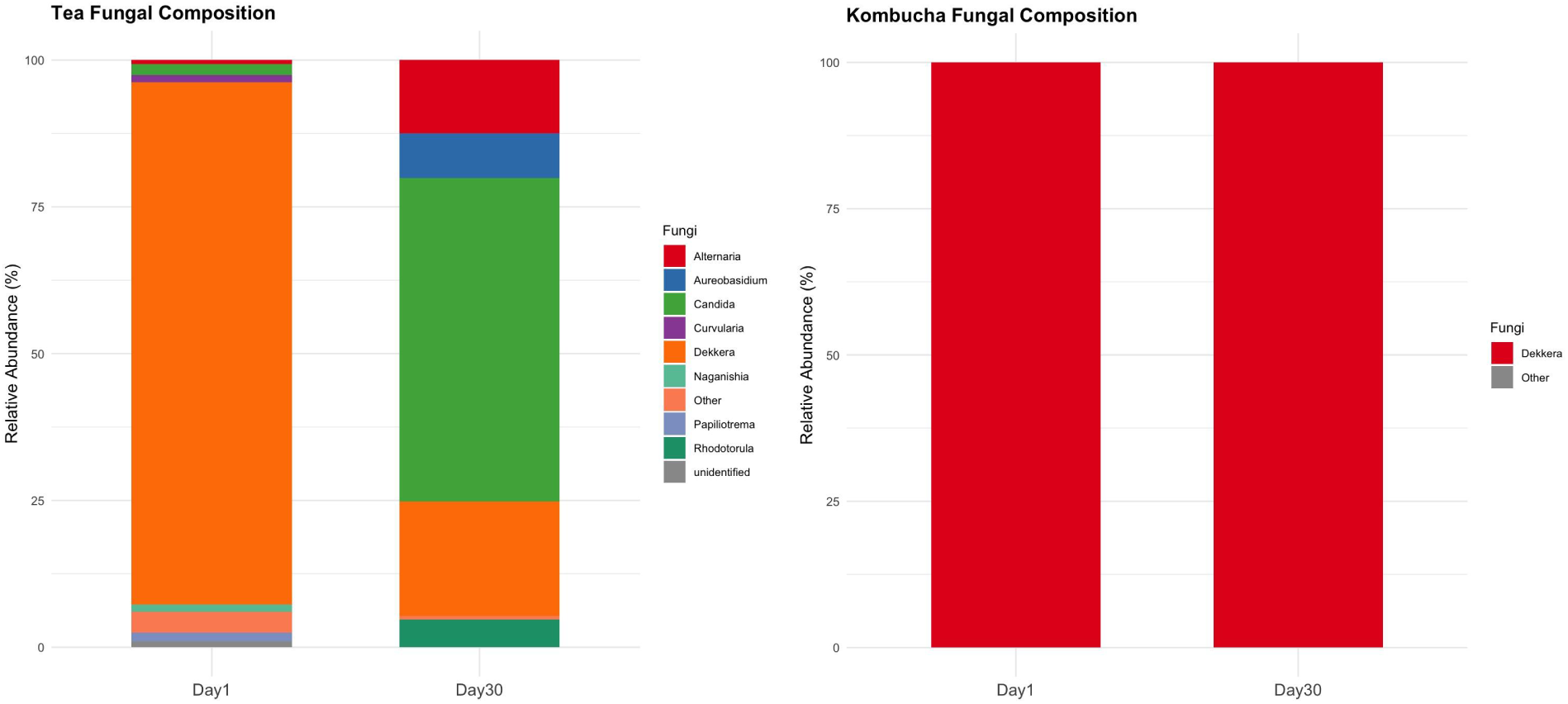
These graphs report the mean relative abundance across all replicates Tea samples changed dramatically through time. *Dekkera* was the most abundant genus on Day 1, but had been reduced to 30% by Day 30. Four other genera of fungi became more abundant, including *Papiliotrema*, and *Alternaria,* with *Candida* becoming the dominant fungus. In kombucha, *Dekkera* accounted for essentially all (99.9%) of fungal read counts, both on Day 1 and Day 30.

## Discussion

### Fermentation may make kombucha and other beverages safer for human consumption

There are no previous studies to our knowledge that explore whether fermented drinks have lower prevalence of human pathogens than comparable non-fermented drinks. We found that microbes from human hand swabs were absent from kombucha’s microbial community after 30 days, while *Burkholderia*, *Staphylococcus*, *Lawsonella*, *Micrococcus*, and *Acinetobacter* were present in the tea samples. This is important because it suggests that the fermentation process of kombucha contributed to making it safer for human consumption during the historical development of the beverage. If kombucha can successfully eliminate and control microbes that we intentionally introduced from people’s hands, we can infer that it is likely equally successful at eliminating the microbes introduced naturally during fermentation processes during small-scale brewing at home or commercially.

### Kombucha has antimicrobial properties that may have prevented invasion

We found that kombucha’s microbial and fungal composition remained relatively stable despite the introduction of foreign microbes from people’s palms. This suggests that kombucha’s microbial composition and the compounds that are created during fermentation may allow it to maintain a diverse and stable microbial ecosystem in the face of perturbations like the introduction of potentially invasive microbes. It is likely that some of the metabolites of bacteria and yeast, including ethanol, and acetic acid contribute to this resilience. These compounds create an environment that can limit the presence of microorganisms not adapted to the membrane-destabilizing effects of low pH and ethanol^11^. For microbes that are not adapted to high acid conditions, such as *E. coli, Helicobacter pylori*, and *Staphylococcus aureus(Micrococcus)*^12^, acidic environments break the plasma membrane and destroy the inner contents of the cell^13^. The antimicrobial properties of this acidic environment likely contributed to the stability of the bacterial and fungal populations in the kombucha samples despite the introduction of potential microbial invaders. Because kombucha is made with 20% kombucha starter, it lowers the pH, which is something that does not occur in the tea. Low pH can suppress the growth of or kill microbes that cannot tolerate acidic conditions.

### The microbial composition of tea changed drastically from Day 1 to Day 30

In contrast to the stability of the kombucha samples, the microbial and fungal composition of the tea replicates in our experiment changed drastically from Day 1 to Day 30, with new bacterial and fungal species increasing in prevalence. These invading microbes were so abundant that they were visible to the naked eye. *Burkholderia* came to dominate tea over the course of the 30-day experiment. One reason for this could be that sugar, a carbon source, supports rapid bacterial growth. The *Burkholderia* genus found in tea has been associated with osteoarthritis and cholangiocarcinoma when present in the human microbiome^14–16^.

### Kombucha’s fungal profile did not change

Kombucha’s fungal composition did not change between Day 1 and Day 30 (Figure 2). *Dekkera anomala* and *Dekkera bruxellensis* were the dominant species in kombucha and maintained their dominance despite the introduction of foreign microbes. *Dekkera* are commonly found in fermented beverages such as beer, wine, and especially kombucha^17,18^. *Dekkera* is known for its ability to produce acetic acid^19^. This may be why other fungal genera did not survive in kombucha. The acetic acid produced by *Dekkera* is a strong antifungal, even against genera such as *Candida*, the dominant fungus in tea samples^20^. *Dekkera’s* acetic acid production and antifungal properties may have prevented the growth of new fungi in kombucha, allowing it to keep its presence and dominance in the face of invasion.

### Kombucha’s SCOBY may protect against invasion

Another mechanism that may have protected the kombucha from invasion is the SCOBY (Symbiotic Community Of Bacteria and Yeast). This SCOBY acts as a physical barrier, potentially protecting the kombucha’s microbial ecosystem by limiting air exposure and hindering the entry of foreign microbes^9^. Future work should investigate the potential protective role of the SCOBY in preventing invasion.

### Future work should investigate diverse mechanisms underlying kombucha’s resilience

Our results suggest that the ecological system of kombucha has the ability to limit invasion from potential pathogens. The products generated through the cooperative fermentation of sugars, for example: acids, ethanol, and cellulose, may be responsible for limiting the growth of invaders in kombucha. It is also possible that yeast and bacteria engage in more direct competitive interactions with invading microbes, such as attacking or physically containing them through the production of microscopic biofilms, in a process that may have parallels with the ways that immune systems function in multicellular organisms. It may also be the case that just by occupying the relevant ecological niches in kombucha, the kombucha microbes prevent invading microbes from flourishing by taking up resources and space. We are investigating these mechanisms of kombucha’s resilience, in particular the potentially mutualistic interactions that help protect the system from invasion, in our ongoing work.

### This study is an example of citizen science

This experiment took place during ASU’s Night of the Open Door event in 2017. Visitors came to ASU to learn about the research going on in the university. Participants approached our booth about kombucha, and we asked them to participate in our study. After they agreed, we swabbed their hands twice and put one swab in tea and the other in kombucha. After participating in the study, participants were encouraged to follow up on our progress by following the lab’s Twitter (now known as X) account @AktipisLab. Lab members posted videos and photos of how the tea and kombucha progressed over 30 days (see Supplementary Fig. 5). We used hashtags including #livingpotion, #citizenscience, and #ASUOpenDoor so participants could easily follow the study’s progress. These results are still available on the platform at the following https://x.com/search?q=%23livingpotion&src=typed_query as of the writing of this paper (https://x.com/search?q=%23livingpotion&src=typed_query). This experiment was repeated at Open Door in the years 2017, 2018, 2019, and 2020. Only the data from 2017 were analyzed and reported here.

## Conclusion

We investigated the impact of introducing foreign microbes from human hands into kombucha and sweet tea. Our key finding is that the microbial and fungal communities within kombucha exhibited remarkable stability, resisting invasion and maintaining their diversity and abundance over 30 days, in stark contrast to the readily invaded sweet tea samples. The sweet tea experienced significant shifts in microbial composition and visual signs of contamination. This resilience in kombucha suggests that the fermentation process fosters a microbial environment that is less hospitable to foreign microbes, potentially contributing to its safety for human consumption. Several factors likely contribute to this protective effect, including the presence of antimicrobial compounds such as ethanol, gluconic acid, and acetic acid produced during fermentation, as well as the physical barrier provided by the SCOBY. Our findings support the hypothesis that fermentation can be a significant factor in enhancing beverage safety by limiting the establishment and proliferation of potentially harmful microorganisms, as demonstrated in our kombucha model system. Further research is underway in our laboratory to elucidate the precise mechanisms and multi-species interactions that underlie kombucha’s resilience to microbial invasion.

## Methods

### Kombucha was prepared for the experiment in the laboratory

The kombucha used in the experiment was brewed and maintained in the Cooperation lab at Arizona State University. Every 10-14 days, the Aktipis lab kombucha culture receives 1 L of sweet black tea. To make the tea for this experiment, we boiled 2 L of deionized water and added 10 grams of Lipton loose-leaf black tea. After steeping for 30 minutes, the leaves were filtered out, and 100 grams of sugar were mixed into the tea. 1 L of sweet tea was reserved for the control condition, and 1 L was used to prepare the kombucha. To make the kombucha, 800 mL of the sweet tea was mixed with 200 mL of kombucha from the source culture in the lab.

### Hand swabs were collected at a citizen science event

We conducted our experiment at Arizona State University’s Open Door event on February 25th, 2017. Conducting this experiment at Open Door allowed us to involve the public in the experiment while educating them about the research we perform in our lab. To set up the experiment, we laid out two large glass containers, where one was filled with 1 L of fresh kombucha (20% starter) and the other was filled with 1 L of sweet black tea. We asked participants to swab their hands with two different sterile swabs and place one in the tea and the other in the kombucha (Supplementary Fig. 4). The swabs were left in their respective containers for 6 hours. At the end of the event, we made four replicates for each condition by transferring them into smaller beakers and leaving them undisturbed in a Biological Safety Level-2 facility for 30 days.

### Kombucha and sweet tea samples were sampled on Day 1 and Day 30

To assess changes in microbial and fungal diversity over time, we analyzed samples from Day 1, immediately after the event, and on Day 30. We began by extracting DNA from the Day 1 and Day 30 samples to send them to be analyzed by Arizona State University’s core facility. Genomic DNA was isolated using the Powerfood Microbial DNA Isolation Kit (MoBio) according to the manufacturer’s protocol. The DNA samples were stored at -20°C and then sent for 16S ribosomal RNA (rRNA) amplicon sequencing and internal transcribed spacer (ITS) amplicon sequencing. Data from 16S sequencing provided us with data on bacterial diversity, and ITS sequencing provided data on fungal diversity (see “Bacterial diversity was captured with 16S sequencing” and “Fungal diversity was captured with ITS sequencing” sections for further details). Negative controls, consisting of two samples each of autoclaved tea and ultrapure water, were processed and sequenced to identify and exclude potential contaminants from the extraction and sequencing steps. 16S and ITS can sometimes identify DNA from things in the extraction buffer and sequencing kits, and we used the negative controls to filter them from the main samples of the experiment. These samples were not part of the main experiment; they were only processed during the extraction setup.

### Bacterial diversity was captured with 16S sequencing

Bacterial community analysis was performed via next-generation sequencing in the MiSeq Illumina platform. Amplicon sequencing of the V4 region of the 16S rRNA gene is performed with the barcoded primer set 515f/806r designed by Caporaso et al. (2012) and following the protocol of the Earth Microbiome Project for the library preparation^21^. PCR amplifications for each sample were done in triplicate, then pooled and quantified using Quant-iT™ PicoGreen® dsDNA Assay Kit (Invitrogen). A no-template control sample was included during the library preparation as a control for extraneous nucleic acid contamination. 240 ng of DNA per sample were pooled and then cleaned using QIA quick PCR purification kit (QIAGEN). The pool was quantified by Illumina Library Quantification Kit ABI Prism^®^ (Kapa Biosystems). Then, the DNA pool was diluted to a final concentration of 4 nM and denatured and diluted to a final concentration of 4 pM with 15% of PhiX. Finally, the DNA library was loaded in the MiSeq Illumina and run using the version 2 module, 2x150 paired-end, following the manufacturer’s directions.

### Fungal diversity was captured with ITS sequencing

Fungal community analysis was performed via next-generation sequencing in the MiSeq Illumina platform. Amplicon sequencing of ITS region was performed with the barcoded primer set ITS1f-ITS2 designed by Smith & Peay 2014 and following the protocol by the Earth Microbiome Project for the library preparation^21^. PCR amplifications for each sample were done in triplicate, then pooled and quantified using Quant-iT™ PicoGreen® dsDNA Assay Kit (Invitrogen). A no-template control sample was included during the library preparation as a control for extraneous nucleic acid contamination. 200 ng of DNA per sample were pooled and then cleaned using QIA quick PCR purification kit (QIAGEN). The pool was quantified by Illumina library Quantification Kit ABI Prism® (Kapa Biosystems). Then, the DNA pool was diluted to a final concentration of 4 nM and denatured and diluted to a final concentration of 4 pM with 25% of PhiX. Finally, the DNA library was loaded in the MiSeq Illumina and run using the version 2 module, 2x250 paired-end, following the manufacturer’s directions.

### Diversity and statistical analyses were performed using QIIME2 and R

Illumina reads were demultiplexed by the sequencing core, after which we used QIIME 2 (v2019.1) for all primary processing. Reads were trimmed and denoised with the DADA2 plugin, chimeras were removed, and amplicon-sequence variants (ASVs) were inferred. Putative contaminants were filtered with decontam in R (prevalence method, threshold = 0.5) using autoclaved tea and water controls. Taxonomy was assigned in QIIME 2 with pre-trained Sklearn classifiers: the SILVA 99 % rRNA database for 16S reads and the UNITE library for ITS reads.

Based on the taxonomic annotation, relative abundances at genus level were measured for both bacterial and fungal data for each sample in QIIME 2.

Alpha-diversity was quantified in QIIME 2 (diversity plugin) as ASV richness—the count of observed amplicon sequence variants—and the Shannon index, which weights those variants by relative abundance. Alpha-diversity metrics were measured post rarefaction based on sequencing read depth from each sample. Metrics were analysed in R (v4.0.3) with Kruskal–Wallis tests (rstatix package) followed, when significant, by Wilcoxon rank-sum pairwise comparisons.

Beta-diversity was evaluated on distance matrices comprising weighted and unweighted UniFrac distances for 16S reads and Bray–Curtis (abundance-based) and Jaccard (presence/absence) distances for ITS reads. UniFrac leverages phylogenetic branch lengths, with the weighted variant further scaling by relative abundance. No established phylogenetic tree knowledge is available for ITS sequences, and thus Bray-Curtis and Jaccard measures were used. Overall community differences were tested with PERMANOVA and, when significant, resolved by pairwise PERMANOVA; ordination plots were rendered with R.

## Data Availability

Raw reads of the samples sequenced are submitted in NCBI SRA archives and available in the accession number PRJNA534131 (https://www.ncbi.nlm.nih.gov/sra/PRJNA534131). SRA archive metadata, QIIME scripts used to analyse the data and R scripts used to remove contaminants (Decontam package), to analyze the stats, plot and visualize data have been made available here: https://github.com/nshrinath1994/NOD_Kombucha_Aktipislab.

## Acknowledgments

We thank Rosa Krajmalnik-Brown, Pamela Winfrey, and Arvind Varsani for their comments and assistance.

This material is based upon work supported by the National Science Foundation under Grant No. 2319438, titled “What rules of life allow collectives to effectively manage risk? Understanding the rules underlying risk management across systems to increase societal resilience.” Any opinions, findings, and conclusions or recommendations expressed in this material are those of the author(s) and do not necessarily reflect the views of the National Science Foundation.

Research reported in this publication was supported in part by the National Cancer Institute of the National Institutes of Health under Award Numbers U54 CA217376, U2C CA233254, R21 CA257980 and R01 CA140657. The content is solely the responsibility of the authors and does not necessarily represent the official views of the National Institutes of Health.

This work was supported by a grant from the Arizona State University’s President Office, which helped establish the Interdisciplinary Cooperation Initiative at ASU. Any opinions, findings, and conclusions or recommendations expressed in this material are those of the author(s) and do not necessarily reflect the views of Arizona State University or its President Office.

This work was supported by a grant from the John Templeton Foundation (grant number 61862, Developing the Field of Cooperation Science: Exploring human nature and cultural evolution through interdisciplinary research). Any opinions, findings, and conclusions or recommendations expressed in this material are those of the author(s) and do not necessarily reflect the views of the John Templeton Foundation.

## Author Contributions

**Gissel Marquez Alcaraz** (Conceptualization, Data curation, Formal Analysis, Investigation, Methodology, Visualization, Writing - Original Draft Preparation, Writing - Review & Editing)

**Shrinath Narayanan** (Conceptualization, Data curation, Formal Analysis, Investigation, Methodology, Visualization, Writing - Original Draft Preparation, Writing - Review & Editing)

**Joe Alcock** (Writing - Review & Editing)

**Jessica D. Ayers** (Investigation/ data collection, Writing - Review & Editing)

**Cristina Baciu** (Data collection, project administration, Writing - Review & Editing)

**Hanna L. Berman** (Writing – Review & Editing)

**Robert R. Dunn** (Conceptualization, Writing – Review & Editing)

**Angelo Fortunato** (Supervision, Methodology, Writing – Review & Editing)

**Alex May** (Conceptualization, Investigation, Methodology, Writing – Review & Editing)

**Carlo Maley** (Conceptualization, Writing – Review & Editing, Funding Acquisition, Supervision)

**Jessica Siegel** (Investigation,Data Curation, Writing – Original Draft Preparation)

**Bryan Yavari** (Writing - Original Draft Preparation, Writing - Review & Editing, Validation)

**Athena Aktipis** (Conceptualization, Investigation, Methodology, Writing – Review & Editing, Validation, Funding Acquisition, Supervision)

## Notes

### Competing Interest Statement

The authors have declared no competing interest.

